# How different immersive environments affect intracortical brain computer interfaces

**DOI:** 10.1101/2024.07.30.605911

**Authors:** Ariana F Tortolani, Nicolas G Kunigk, Anton R Sobinov, Michael L Boninger, Sliman J Bensmaia, Jennifer L Collinger, Nicholas G Hatsopoulos, John E Downey

## Abstract

As brain-computer interface (BCI) research advances, many new applications are being developed. Tasks can be performed in different environments, and whether a BCI user can switch environments seamlessly will influence the ultimate utility of a clinical device. Here we investigate the importance of the immersiveness of the virtual environment used to train BCI decoders on the resulting decoder and its generalizability between environments. Two participants who had intracortical electrodes implanted in their precentral gyrus used a BCI to control a virtual arm, either viewed immersively through virtual reality goggles or at a distance on a flat television monitor. Each participant performed better with a decoder trained and tested in the environment they had used the most prior to the study, one for each environment type. The neural tuning to the desired movement was minimally influenced by the immersiveness of the environment. Finally, in further testing with one of the participants, we found that decoders trained in one environment generalized well to the other environment, but the order in which the environments were experienced within a session mattered. Overall, experience with an environment was more influential on performance than the immersiveness of the environment, but BCI performance generalized well after accounting for experience.

## Introduction

Brain-computer interfaces (BCIs) hold the promise of improved quality of life for many people with chronic motor impairments. Intracortical BCIs can provide people with cervical spinal cord injury, brainstem stroke, or ALS with the abilities to communicate via decoded cursor control [1,2], handwriting [3], or speech [4]; to control the movements of a virtual [5,6] or real robotic arms [7–12]; or, through functional electrical stimulation, to move their paralyzed hand [13,14] and arm [15]. For BCI users to accomplish a variety of tasks, their abilities will need to generalize across environments.

Virtual reality (VR) technologies enable researchers to create virtual environments that feel more natural and immersive for BCI users, and can be used to train BCI decoders that are used with physical robots or functional electrical stimulation [6,15]. Visual feedback is an important element to consider as modifying the design of a virtual limb [16,17] or its alignment relative to a user’s native limb [18] can impact overall performance. Further, manipulating the relationship between the BCI users intended movement and action performed by the virtual limb decreases an individual’s sense of agency and embodiment of that limb [17,19]. These findings highlight the importance of clear and accurate visual feedback. Researchers have begun testing how different setups, such as a VR headset or a standard monitor display, effect BCI usability; however, results have varied from improvements when working in VR [20], to mixed effects [21], or no difference between the two conditions [22]. These studies have primarily relied on non-invasive or low-dimensional BCIs which can work with an abstracted strategy to control movement; little is known about how the immersiveness of the environment will affect more intuitively controlled higher degree of freedom BCIs.

Here we began by changing the immersiveness of the environment within which two BCI users controlled a virtual robotic arm. They either wore a VR headset, providing a fully immersive experience with the arm, or viewed the arm on a television (TV) at a distance, providing an abstracted experience with the arm. We asked whether the immersive condition made it easier for the participants to control the arm. We found that both participants performed slightly better in the environment in which they had routinely performed BCI control before the study, for one participant that was the VR environment and for the other that was the TV environment. Indeed, we further analyzed the underlying neural data and found that multi-unit activity recorded from the precentral gyrus maintained similar tuning regardless of which environment the participant’s robotic prosthetic was viewed in. Finally, to investigate the generalizability of decoders, one of the participants attempted to use decoders trained in each environment to complete tasks in both environments. He succeeded with decoders trained in the VR environment regardless of which environment.

Overall, we found that BCI control typically generalized between environments. However, there were some differences that were presumably related to previous experience with the condition in both the short and long terms, which should be considered when developing generalizable decoders.

## Methods

### Participants

Both participants enrolled in a multi-site clinical trial of an intracortical sensorimotor BCI (registered on clinicaltrials.gov, NCT01894802) that was conducted under an FDA Investigational Device Exemption and provided informed consent prior to any experimental procedures. All procedures were approved by the Institutional Review Boards at the University of Pittsburgh or the University of Chicago. Participant C1 (male), 57 years old at the time of implant, presented with a C4-level ASIA D spinal cord injury (SCI) that occurred 35 years prior. Participant P4 (male), 30 years old at the time of implant, presented with a C4 ASIA A SCI that occurred 11 years prior. C1 completed this study between 645 and 989 days after the electrode arrays were implanted. P4 completed this study between 251 and 385 days after the electrode arrays were implanted.

### Neural Recording Setup

Both participants had two microelectrode arrays with 96 electrodes each (Utah Arrays, Blackrock Neurotech, Salt Lake City, UT) implanted in the precentral gyrus. Signals from these electrodes were recorded using the NeuroPort system at 30 kHz, high-pass filtered with a 1^st^ order 750 Hz filter [23], and every crossing of a voltage threshold (−4.25 RMS, set at the start of each recording session) was counted in 20 ms bins and used for offline analysis and online decoding.

### Grasp and Transport Task

Participants performed a 4 degree of freedom grasp and transport motor imagery task in one of two environments. In this task, participants attempted to reach an object in 3-dimensional space, grasp the object, transport the object to a new target location, and then release the object at that site. Once all four phases were completed a new trial started immediately with the appearance of a new target object. The phase was completed successfully when the hand was within 5 cm of the target and the hand opened or closed (within 20% of range of motion) as instructed. If any of the phases were not completed within 10 seconds, the trial was failed and the hand automatically returned to the center home position before a new trial began. We asked participants to complete this task in two different environmental conditions, either in a three dimensional VR headset (Figure 1a, participant C1: Index, Valve Corp, Bellevue, WA USA, participant P4: Quest, Meta, Menlo Park, CA, USA) or with a 2 dimensional TV screen (Figure 1b, participant C1: TV screen 100 × 85 cm at a distance of 244 cm, participant P4: PC monitor 27.5 × 15 cm at a distance of 70 cm), within the same experimental session. All trials for a given condition were completed before switching to the other environmental condition and the order was balanced across experimental sessions (participant C1: 3 sessions, participant P4: 2 sessions).

**Figure 1.**
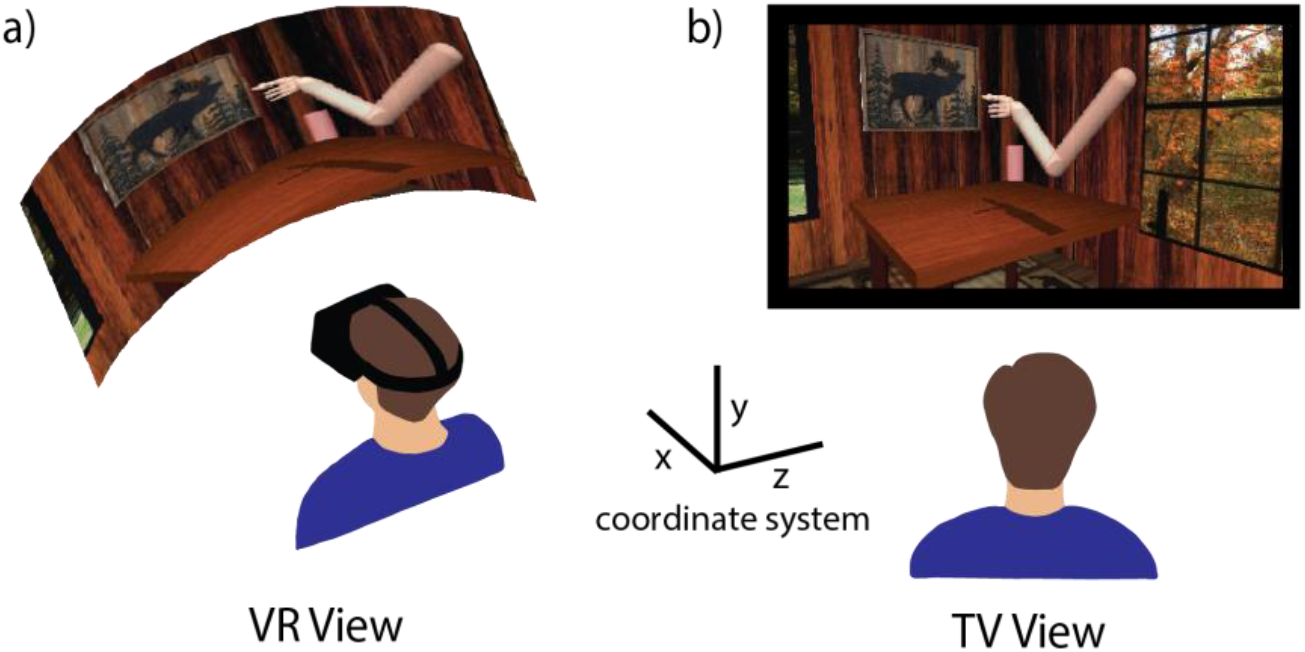
Experimental Setup. **a)** In the VR experimental condition, the participant wears a virtual reality headset where the field of view moves to match their head movements to interact with the environment. **b)** In the TV experimental condition, the participant looks at a fixed view on a screen. Inset displays coordinate system used in the MuJoCo virtual environment.

### Decoder Training

Each environmental condition started with initial calibration sets performed in the respective environment, where the participant was instructed to observe and imagine they are completing the task while the computer controlled the actual movements of the hand (participant C1: 54 trials, participant P4: 36 trials). Using these observation sets we trained an optimal linear estimator decoder [6] which the participants used to complete an error-limited calibration step. During this step the participants attempted to control the arm but the computer prevented any deviations in the path from the start position to the target location by eliminating velocities orthogonal to that path (participant C1: 54 trials, participant P4: 36 trials). These error-limited sets were used to train another decoder which the participants then used to perform unassisted BCI-controlled trials of the task. Participant C1 attempted 171 trials in each condition across 3 experimental sessions, while participant P4 completed 108 trials in each condition across 2 experimental sessions. During the first two calibration steps of the decoder training, participant C1 performed a 5 degree of freedom version of the task, rotating the wrist before grasping targets, but that was not done during BCI-controlled trials. For participant P4, during the BCI-controlled trials he only had control of the type of movement needed to reach the current target (i.e. no grasp movements were permitted while reaching to the target, and no arm movements were permitted while grasping the target).

### Performance Metric Calculations

To quantify BCI performance we analyzed several metrics: failure rates, completion times, and normalized path length. As this task is performed in discrete phases, we can segment each of these analyses to look at the reach and transport phases individually. The failure rate is defined as the percentage of trials where the participant failed to reach the target in the allotted time (10 seconds). Completion time is a measure of how quickly the participant was able to successfully move to the target location. If a phase was failed, then the failure time (10 seconds) was used in the analysis, though the results were still significant if failed phases were excluded from the analysis. Finally, normalized path length is the ratio of the of the length of the executed path to the minimum distance from the start point to the outer edge of the target was calculated both in total and for each translation dimension individually (x, y, and z). For the individual dimension analysis, we normalized the distance the participant traveled in a particular dimension by the maximum target separation distance allowed in that dimension (x=0.17, y = 0.42, z = 0.5). A normalized path length of 1 indicates the participant traversed the ideal path to reach the target. Phases where the new target location appeared in the same location as the previous target were excluded from analysis.

### Neural Tuning

To quantify neural tuning, we calculated the preferred direction for each active channel for each environmental condition. We defined an active channel as having an average firing rate above 3.5 Hz during the task, to ensure enough modulation to stably calculate tuning, giving us 279 active channels across 3 datasets for participant C1, and 336 active channels across 2 datasets for participant P4. Some neurons may have been measured multiple times in different sessions, but overlap was likely low given the amount of time between sessions [24]. Preferred directions are defined by fitting a linear encoding model for each channel as follows:

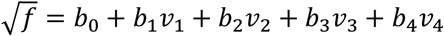

where f is the firing rate on a given channel and v was the velocity of movement for a given degree of freedom. Fitting the b coefficients through linear regression using data from full trials, not individual phases, defines the tuning of a channel. The preferred direction of the channel was defined by the angle vector of the first three velocity coefficients. The depth of modulation was calculated as the magnitude of the same vector. We parsed the data in half for each condition and calculated a preferred direction for each channel in both halves of the VR and TV conditions. We could then calculate the shift in each channel’s preferred direction within condition (e.g., VR to VR), across conditions (VR to TV), or across conditions where the channel identity had been shuffled to determine how stable the tuning properties are across conditions. We also calculated the percentage change in depth of modulation for each unit between tasks.

### Online Decoder Comparison

To test how well a decoder trained in one condition generalizes across conditions, we trained two separate decoders (one for the VR environment and one for the TV environment) at the start of an experimental session and then tested both of these decoders in both environments on the same grasp and transport task with participant C1. For a given day, we would start working in the VR environment and train a decoder (VR decoder) according to the methods detailed above. The participant was then allowed 2 short unassisted sets to practice using the decoder in that environment to ensure reasonable performance of the decoder for the task. We then immediately switched to the TV environment and repeated this process, training a new decoder (TV decoder). After satisfactory training of both decoders was completed, we switched back to the VR environment and began testing trials. Testing trials were completed in blocks of 18 trials at a time using either the VR decoder or TV decoder. After finishing all testing trials in the VR environment, we switched to the TV environment and again completed testing trials with alternating decoder blocks (Figure 2). The participant was not aware which decoder was active during testing trials. We collected 4 sessions in total, 2 starting in the VR environment and 2 starting in the TV environment. For analysis, failure rates were calculated using data across all 4 testing sessions while completion times and normalized path length were calculated using data from only the 2 sessions where training started in the VR environment due to the excessive failure rate of the TV decoder in the excluded sessions.

**Figure 2.**
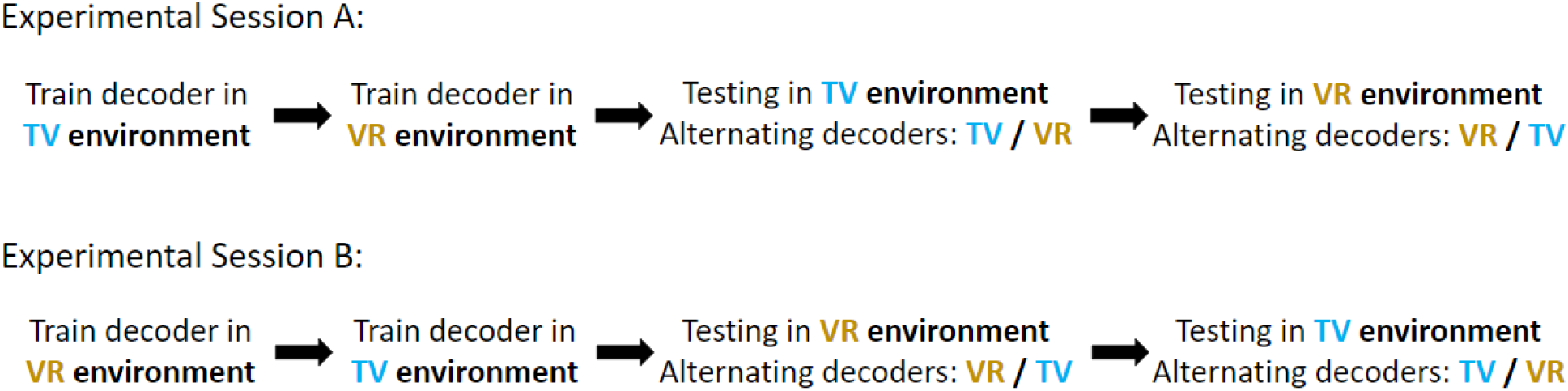
Online decoder training paradigm. To test decoder generalizability with participant C1, for a given experimental session, we fully trained separate decoders in the TV environment and VR environment and then alternated testing both decoders in both environments. The starting environment was balanced across sessions.

## Results

The participants each completed multiple sessions comparing their ability to control an arm in a virtual environment, either projected on a TV or in VR (Figure 1). Throughout this task, participant C1 noted a strong preference for the VR condition, as working in that environment felt much more natural for him. Participant P4 noted that although the task was easier for him in the TV condition because he didn’t have to move his head at all to see the entire environment, he felt closer to the workspace in the VR environment and more like the virtual arm was his own.

### Failure Rates

In line with the subjective reports from participants, C1 was more successful in the VR condition and P4 performed better in the TV condition. Across 3 test sessions, during unassisted use of the optimal linear estimator decoder, participant C1 failed fewer trials in the VR condition than in the TV condition (26.9% and 38% respectively, p = 0.0282, chi-squared test). He was slightly less likely to fail to reach the object successfully while working in the VR environment as opposed to the TV environment (9.4% vs 14.1% failure rate, p = 0.2, chi-squared test) and much less likely to fail to transport the object (4% vs 18.5% failure rate, p < 0.001, chi-squared test). However, participant P4 failed fewer trials in the TV condition than in the VR condition (0% and 24.1% respectively, p < 0.001, chi-squared test). Accordingly, he only failed during reach and transport in the VR condition (12.9% and 9.8% for reach and transport, respectively, p < 0.001 and p < 0.01, chi-squared test, Figure 3a).

**Figure 3.**
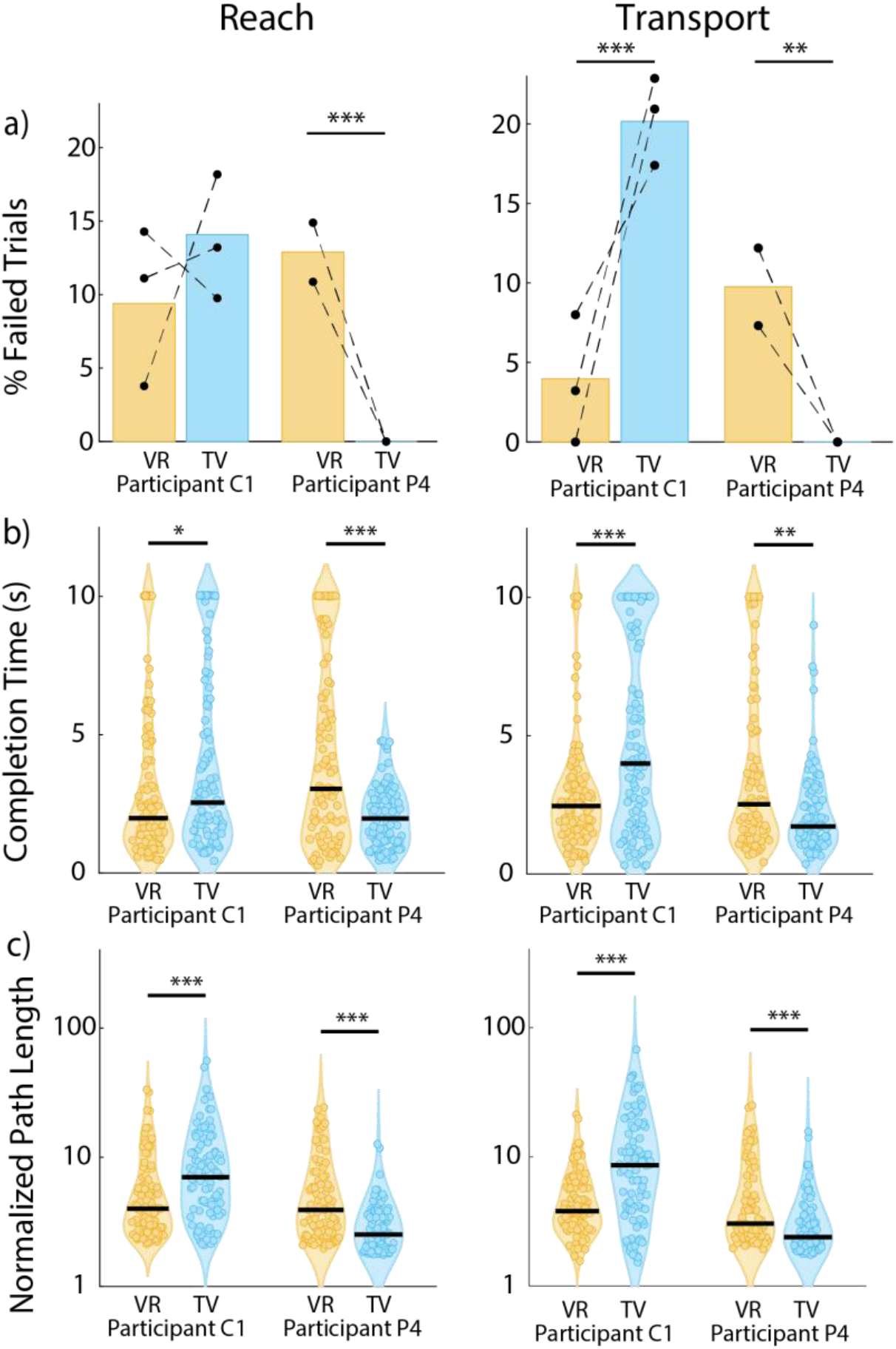
Performance metrics for VR versus TV experimental conditions. **a)** Failure rates for participants C1 and P4 during the reach and transport phases. Data collected during the same experimental session are connected via a dotted line. **b)** completion times, and **c) normalized** path length from the reach (left column) or transport (right column) phases. Black lines indicate median values. Path length is plotted on a log scale. *p<0.05, **p<0.01, *** p<0.001, chi-squared test for failure rates, Wilcoxon rank-sum test for completion times and path length.

**Figure 4.**
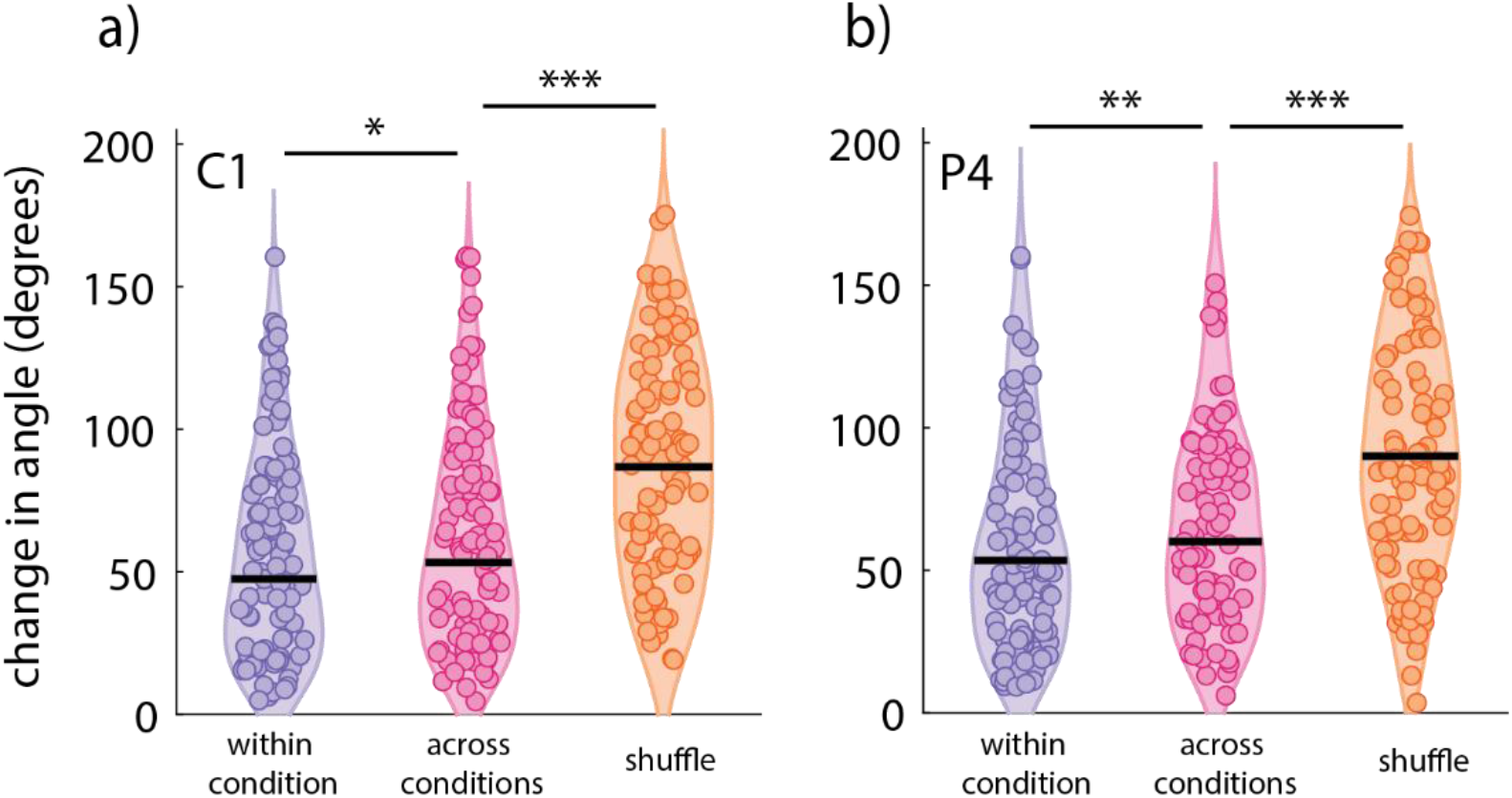
Neural tuning across conditions. Shift in preferred direction angle for **a)** participant C1 and **b)** participant P4. Within condition quantifies changes in tuning within the same environment pooled across VR and TV. Across condition describes changes in tuning between the VR and TV environments. The shuffle condition is a comparison of changes in across condition tuning where the recording channel numbers have been shuffled. Individual points are a singular channel (279 channels for participant C1, 336 channels for participant P4). Black lines indicate median values. *p<0.05, **p<0.01, *** p<0.001, K-S test.

### Completion Times

Completion times were dependent on the testing environment, again matching the reported preference of each participant. Participant C1 took 0.56 seconds less to complete the reach phase in the VR condition (medians: 1.98 sec in VR, 2.54 sec in TV, p = 0.0457, Wilcoxon rank-sum test), and 1.55 seconds less to complete the transport phase in the VR condition (medians: 2.46 sec in VR, 3.99 sec in TV, p<0.001, Wilcoxon rank-sum test). Inversely, P4 took 1.07 seconds less to complete the reach phase in the TV condition on average (medians: 3.04 sec in VR, 1.97 sec in TV, p<0.001, Wilcoxon rank-sum test). He also took 0.80 seconds less to complete the transport phase in the TV condition (medians: 2.52 sec in VR, 1.72 sec in TV, p<0.01, Wilcoxon rank-sum test, Figure 3b).

### Normalized Path Length

We found that C1 made shorter, that is more direct, movements to the target in the VR condition for both the reach and the transport phases of the task (4x minimum in VR, 7x minimum in TV for reach, p<0.001 and 3.8x minimum in VR, 8.6x minimum in TV for transport, p<0.001, Wilcoxon rank-sum test). Conversely, P4 made shorter reaches in the TV condition in both phases (3.9x minimum in VR, 2.52x minimum in TV, p<0.001, for reach and 3.04x minimum in VR, 2.39x minimum in TV, p<0.001, for transport, Wilcoxon rank-sum test, Figure 3c). Normalized path lengths were analyzed separately for each reach dimension to assess whether difficulties with depth perception led to differences in performance across dimensions, but no such pattern appeared (Figure S1).

### Stable unit tuning across conditions

After quantifying the differences in performance across both subjects, we were interested in how similar or different neural activity was between the conditions. To do this, we split the data from each condition on each day into two parts and calculated the shift in the preferred direction for each active channel between the two partitions within and across conditions. The shift within the same condition was used to establish the baseline for the stability of the tuning. We found that there was a slightly larger shift in the preferred direction across conditions than within conditions for both participants (53.3° vs 47.4°, p<0.05, for participant C1, and 60.2° vs 53.4°, p<0.01, for participant P4, Wilcoxon rank-sum test). To compare this shift to chance we shuffled the identities of the channels and recalculated the change in preferred direction across conditions with the shuffled channel labels. We found that tuning was significantly different across conditions with the channel identities shuffled compared to the non-shuffled across condition comparison in both participants (86.8° vs 53.3°, p<0.001, for participant C1, 90.1° vs 60.2°, p<0.001, for participant P4, Wilcoxon rank-sum test, **Error! Reference source not found**.). C1 showed no significant difference in depth of modulation between environments (median 8.2% higher in VR, p = 0.46, KS Test) while P4 showed higher depths of modulation in the TV environment (median 36.6% higher in TV, p = 0.0007, KS Test, Supplementary Figure 2). This demonstrates that while it is difficult to precisely calculate the preferred direction on this data, they were much more stable across conditions than would be expected by chance, while the depth of modulation was higher in the preferred environment for P4 but not C1.

### Decoder comparison testing

While performance was better in the condition that each subject was most practiced in, and neural tuning did not show meaningful differences between conditions, we were interested in whether decoders trained in one condition would yield similar control in the other condition. To determine how transferable decoders were, C1 completed 2 sessions where a decoder was first trained in the VR environment, and then a second decoder was trained in the TV environment. Both decoders were then tested in both environments in alternating blocks. The same experiment was repeated in 2 other sessions with the order of decoder training switched.

Out of all conditions, participant C1 performed best in the VR environment when using the VR decoder. We found that when training started in the VR environment followed by the TV environment, both decoders performed similarly when testing in either environment. Failure rates were similar for both the VR and TV decoders in the VR environment (2.1% vs 6.3%, p=0.2, chi-squared test) and in the TV environment (4.2% vs 2%, p=0.4, chi-squared test) during the reach phase, as well as the transport phase (8% vs 1.2% in VR environment, p = 0.0299 chi-squared test; 7.7% vs 6.7% in TV environment, p=0.7, chi-squared test, Figure 5a).

**Figure 5.**
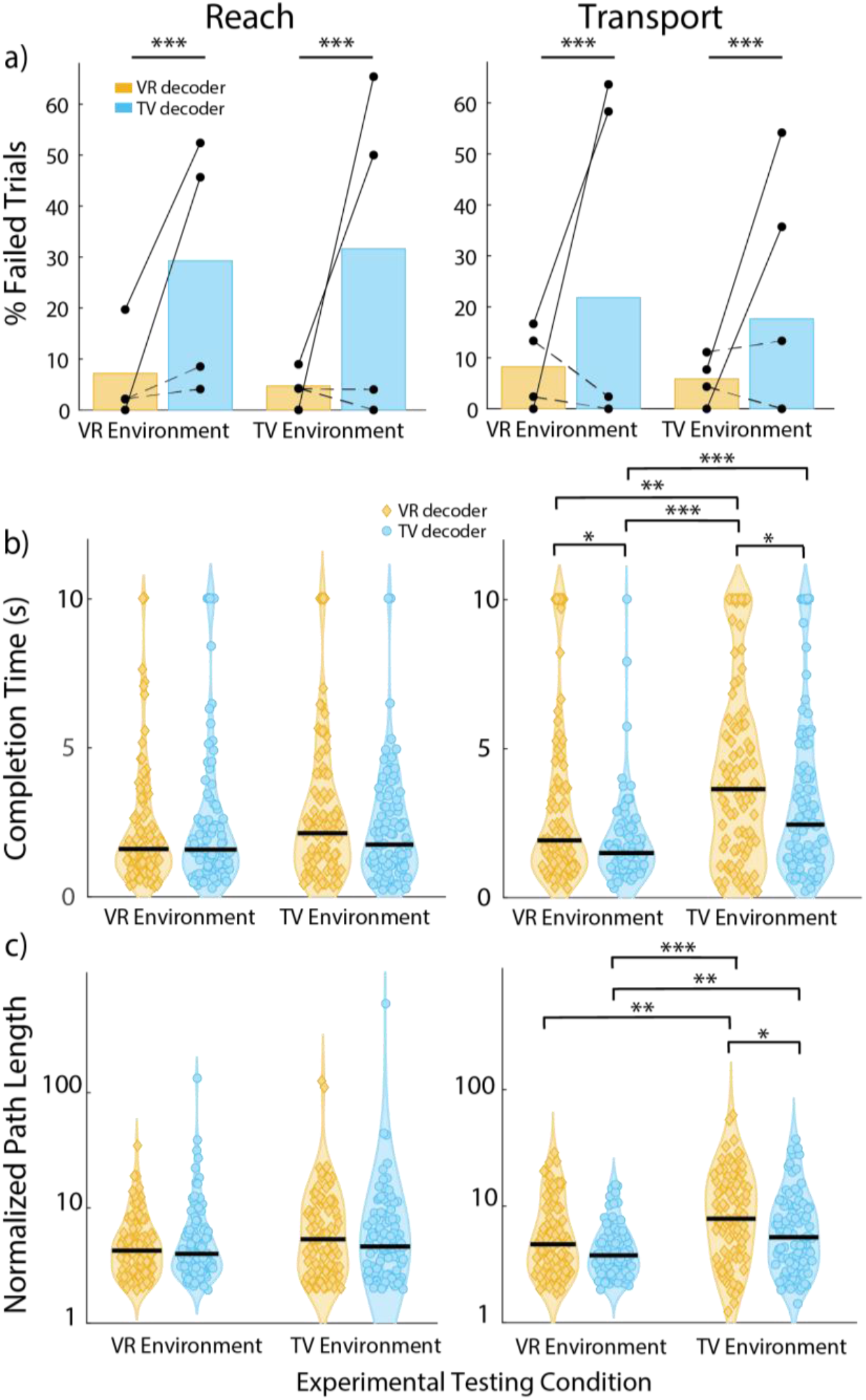
Decoder comparison performance metrics. **a)** Failure rates during online decoder comparison testing for reach and transport phases. Yellow indicates performance when using the decoder trained in the VR environment. Blue indicates performance when using the decoder trained in the TV environment. Solid lines indicate experimental testing sessions where training started in the TV environment. Dashed lines indicate experimental testing sessions where training started in the VR environment. **b)** completion times, and **c)** normalized path length during online decoder comparison testing for the reach (left column) or transport (right column) phases for sessions where training started in the VR environment. Black lines indicate median values. This task was only completed by C1. Path length is plotted on a log scale. *p<0.05, **p<0.01, *** p<0.001, chi-squared test for failure rates, Wilcoxon rank-sum test for completion times and path length.

However, the order of decoder training had a strong impact on performance in different environments. When the TV decoder was trained first, it failed to transfer to the VR environment, even though the VR decoder still successfully transferred to the TV environment. Using the TV decoder in the VR environment, participant C1 failed to reach to the object far more frequently than when using the VR decoder (49% vs 9.8%, p<0.001, chi-squared test). This was similar for the TV environment (57.7% vs 4.5%, p<0.001, chi-squared test), and for the transport phase in the VR (61% vs 8.3%, p<0.001, chi-squared test) and TV (45% vs 3.8%, p<0.001, chi-squared test) environments (Figure 5a). This unidirectional decoder transfer performance (i.e., VR decoders always work in VR or TV, but TV decoders only work if trained after the VR decoder) indicates that there are some differences in the environments that need to be accounted for.

The exceptionally poor performance of TV decoders trained first was apparent from the failure analysis and did not require further analysis of movement time or distance. Further analysis was warranted for the two decoders when they both had low failure rates (i.e., when the VR decoder was trained first). The participant performed about equally with both decoders across both environments during the reaching phase of the task (Figure 5b&c) suggesting that decoders can be used successfully in different environments than they are trained in. However, on average he was able to transport the object to the target location faster when working in the VR environment than in the TV environment with both the VR decoder (1.92 sec in VR, 3.64 sec in TV, p<0.01, Wilcoxon rank-sum test) and the TV decoder (1.50 sec in VR, 2.45 sec in TV, p<0.001, Wilcoxon rank-sum test). Additionally, he made more efficient movements when working in the VR environment with both the VR decoder (4.69x minimum in VR, 7.78x minimum in TV, p<0.01, Wilcoxon rank-sum test) and the TV decoder (3.78x minimum in VR, 5.41x minimum in TV, p<0.01, Wilcoxon rank-sum test).

## Discussion

### Experience is more important than immersiveness

Here we show that participants were able to complete a grasp and transport task, carrying out the same movements in a virtual environment but viewed either as a two-dimensional projection on a TV screen or with binocular depth cues in a VR headset. Both participants performed better with the view that they had primarily used in previous work. Participant C1 had almost exclusively performed BCI tasks with the VR headset for the 2 years since starting the study (e.g. [5]). Participant P4 had only performed BCI tasks on the TV in the 8 months since starting the study. While there were significant differences in performance in the different conditions, the magnitude of the differences was relatively small and both participants were able to complete the tasks satisfactorily in both conditions. This result adds an interesting piece of information to previous work that showed conflicting results as to how a VR environment could influence BCI control [20–22]. It may be that the importance of immersiveness is small or that the importance of experience in an environment is large, but in either case a device for long-term use should be able to utilize either type of environment.

### Stable neural tuning

The tuning of neurons in motor cortex to desired movement showed changes in preferred directions between environmental conditions, that were only slightly larger than between partitions within one environment, and significantly less than chance levels. One participant showed stronger tuning (i.e. larger depth of modulation) in his preferred environment, but the other did not. This is consistent with the relatively minor differences in BCI control between the two environmental conditions but was not a given since changes in perspective of the arm could have resembled changes that required reaiming in non-human primate BCI experiments [25]. It may be that showing the same arm from different views allowed the participants to embody the movements in a way that non-human primates couldn’t when completing cursor control tasks. Future work can explore the differences in robustness of tuning to attempted arm control compared to cursor control.

### Decoder generalizability

Beyond testing whether the participants could achieve comparable control in both environments, we wanted to test whether a decoder trained in one environment generalized to the other. Generalizability of decoders will be an important feature for clinical BCIs that will need to be usable in all the various conditions that occur in a user’s life to maximize utility. Previously we found that decoders trained using movements in a VR environment on a 3D TV worked well for a physical robotic arm, but had been unable to make a precise comparison because of differences in how tasks were performed in the real world and the VR environment [6,12]. Here we found that for a user who usually trained in VR, decoders trained in VR always generalized to the TV condition, but the decoders trained in the TV condition at the start of a session failed to maintain performance in either environment once a VR decoder was trained. This result implies that there are behavioral or strategic factors during training that can influence generalizability. To better understand this phenomenon, further study with more subjects and a wider variety of environments will be required.

Taken together, all of these results provide a picture of BCI performance which is not critically dependent on the environment in which it occurs. There are small changes based on environment, likely related to the participants’ level of experience in each environment. Decoders trained in one environment can readily generalize to the other environment under certain circumstances. We expect that this relative lack of dependence on environment will lead to innovative ways of providing training that enable broadly beneficial BCI assistive devices for patients with a variety of needs.

## Acknowledgments

This work was supported by the National Institute of Neurological Disorders and Stroke, UH3 NS107714, R01 NS130302, and T32 NS121763.

AFT, SJB, NGH and JED conceived the study. ARS developed the virtual environment. AFT, ARS and JED developed the experiments. AFT and NGK ran experiments and collected data. AFT and JED analyzed data. MLB, SJB, JLC, NGH and JED acquired funding and provided supervision. AFT and JED drafted manuscript. AFT, NGK, ARS, MLB, JLC, NGH and JED reviewed and edited the manuscript.

## Supplementary information

**Supplemental Figure 1.**
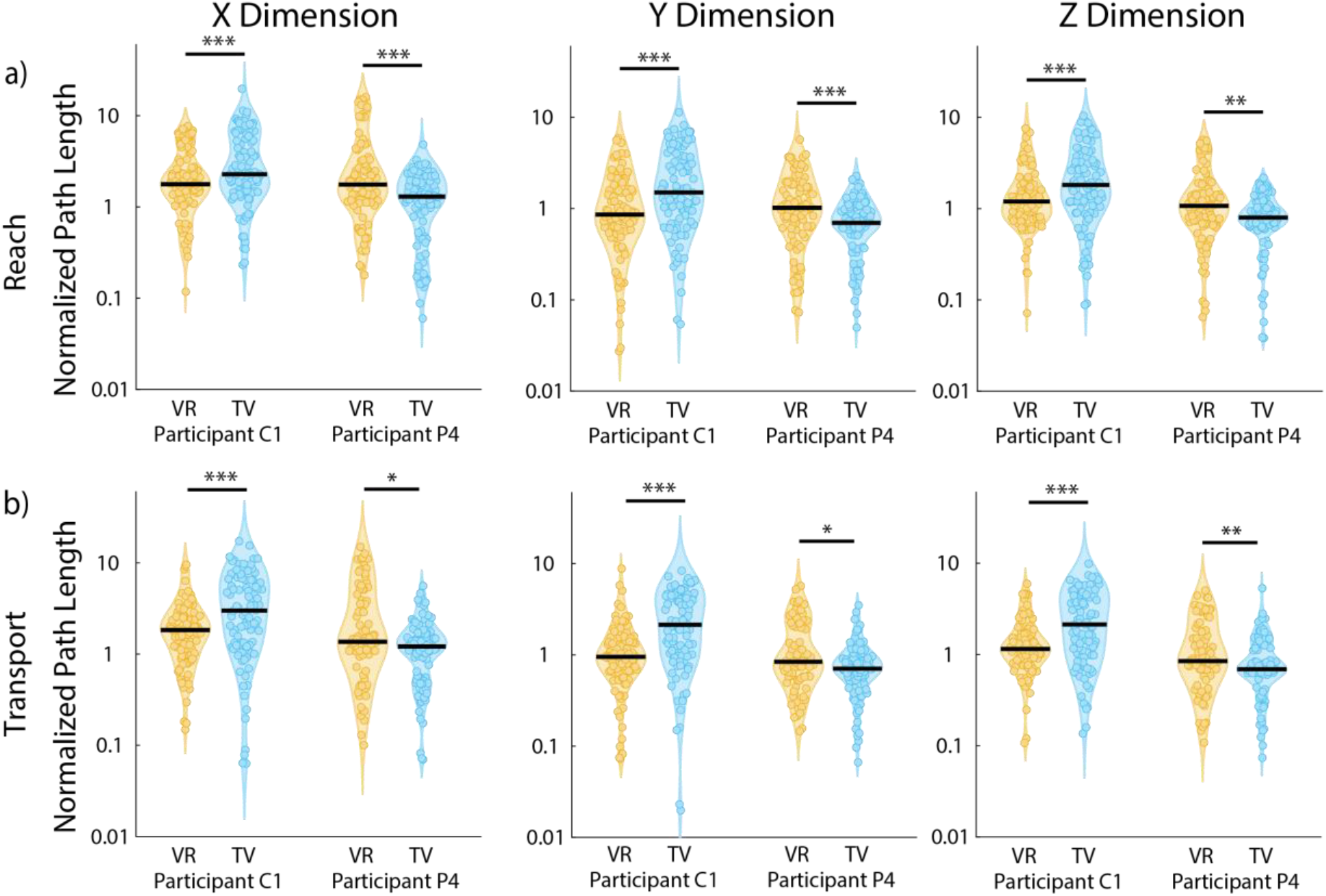
Normalized path length by dimension of translation. Normalized path length for individual dimensions of translation for participants C1 and P4 during control in either the VR or TV condition for both the **a)** reach and **b)** transport phases. X is front/back, Y is up/down, Z is left/right. Black lines indicate median values. *p<0.05, **p<0.01, *** p<0.001, Wilcoxon rank-sum test.

**Supplemental Figure 2.**
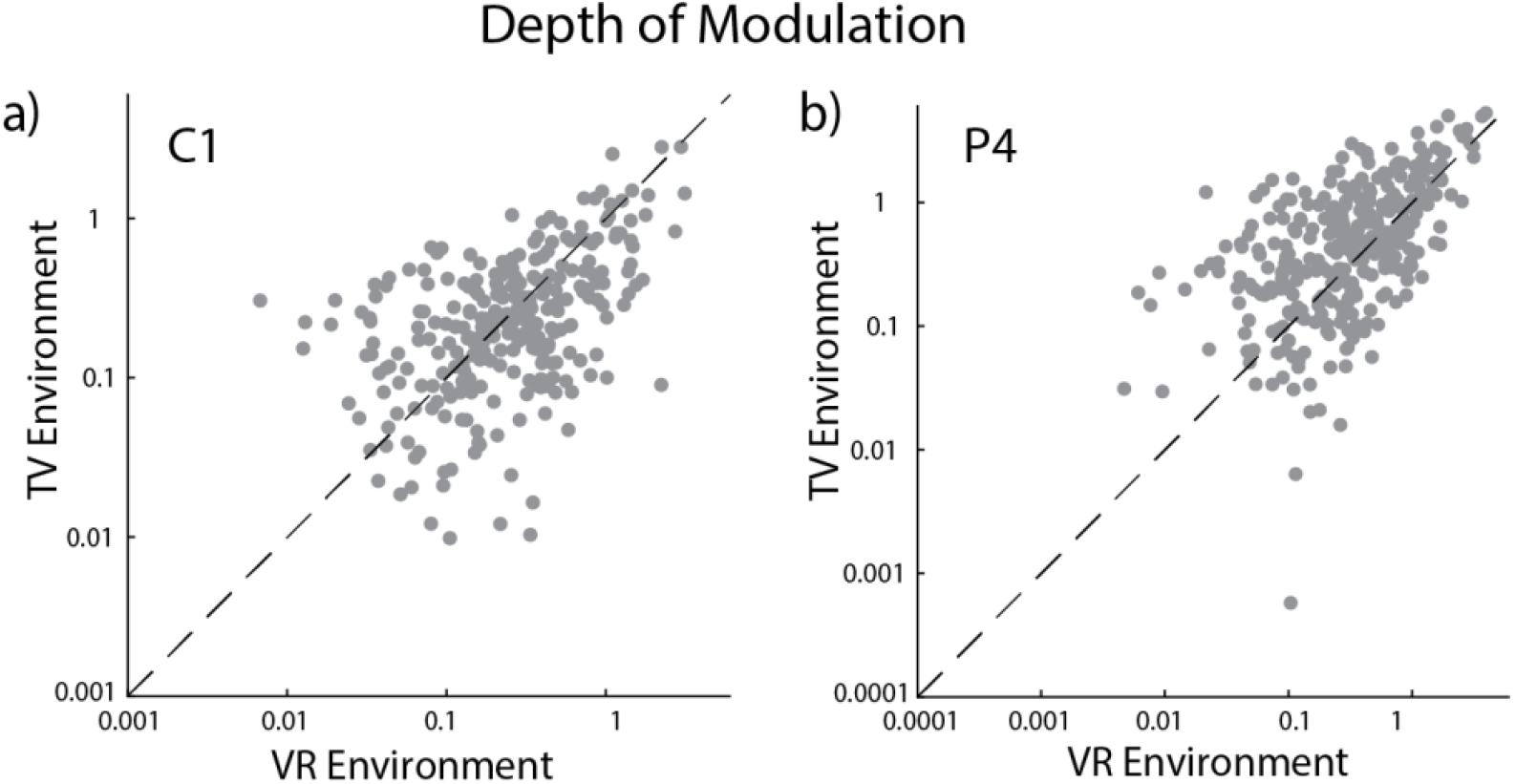
Depth of modulation for each active channel. Depth of modulation for each active channel for **a)** participant C1 and **b)** participant P4 calculated during observation trials. Data plotted on a log-log scale.

